# Overflow of Working Memory from an Easily Accessible Active State

**DOI:** 10.1101/2022.11.02.514822

**Authors:** Takuya Ideriha, Junichi Ushiyama

**Author notes:** Correspondence (J.U.), (T.I.). **Author contributions** Conceptualization, T.I. and J.U.; Methodology, T.I.; Software Development, T.I.; Data acquisition, T.I.; Data Analyses, T.I.; Visualization, T.I.; Interpretation, T.I. and J.U.; Writing – Original Draft, T.I.; Writing – Review & Editing, T.I. and J.U.; Supervision, J.U.; Funding Acquisition, T.I. and J.U.

## Abstract

Working memory is a system that realizes short-term memory retention and is essential to everyday activities. Recently, it has been suggested that working memory items can be maintained in both an active state with sustained neural activity and a latent state without any explicit neural activity. However, how easily we can retrieve memories in each state and how much information can be retained in the active state remains unknown. Here, we addressed these questions by adopting a novel experimental paradigm for measuring the reaction time required to recall a letter of the alphabet when presented with a corresponding color, based on memorized color/letter pairs. The results demonstrated that when participants retained only two pairs of items, they could recall both pairs similarly quickly. However, when there were two or more memorized pairs, the most recent or internally attended pairs were recalled more quickly than the other pairs. These results suggest that memories in the “active” state are accessed first and memories that have overflowed into the “latent” state are accessed afterward. Additionally, the capacity of the easily accessible active state is extremely limited (to approximately two chunks of information) compared with the traditionally considered working memory capacity (four to seven chunks). These results provide robust behavioral support for the co-existence of multiple states in working memory and underscore the need for a more detailed classification of the working memory capacity.

**SIGNIFICANCE STATEMENT:** We sometimes cannot quickly retrieve what we are sure we remember. Regarding working memory retrieval, we observed a phenomenon that the most recently encoded memory was retrieved quickly whereas others were retrieved afterward, showing an “overflow” of memory items from an easily accessible active state. This observation has two critical implications: 1) when maintaining multiple items in working memory, not all items are functionally the same, and there is a dissociation between easily accessible active items and items that have overflowed to a latent state; 2) the number of items that can be maintained in the active state is two, which is remarkably smaller than the traditionally considered working memory capacity of four, necessitating the update of working memory models.

## INTRODUCTION

Working memory is the short-term memory system that provides a basis for our everyday behavior, from daily activities such as cooking to complex reasoning (Baddeley & Hitch, 1974). In the history of psychology and neuroscience, working memory items have long been implicitly assumed to be maintained in a single state. However, this assumption has recently been challenged and it has been suggested that there are several ways to maintain working memory items (Kamiński & Rutishauser, 2020; Stokes et al., 2020).

Supporting this view, several seemingly conflicting models of the maintenance of working memory items have been proposed. Traditionally, working memory items have been considered to be maintained in an “active” state by the sustained firings of neurons in the prefrontal cortex (Funahashi et al., 1989; Fuster & Alexander, 1971). Opposing this model, recent studies have demonstrated that working memory information can be maintained in a “latent” state without sustained firings (Rose et al., 2016; Spaak et al., 2017; Stokes, 2015). For instance, Spaak et al. (2017) showed that sustained firings disappeared under certain circumstances, even though participants could successfully recall the memorized item afterward. A suggested mechanism of the latent state is short-term synaptic plasticity (Mongillo et al., 2008; Zucker & Regehr, 2002). Considering these conflicting reports, it is possible that working memory items can be maintained in several states (Kamiński & Rutishauser, 2020; Stokes et al., 2020).

Furthermore, recent evidence supports the co-existence of active and latent states in parallel in working memory. When humans hold multiple items concurrently in memory and attend to one of them, the attended item has been reported to be accurately decodable through functional magnetic resonance imaging (Lewis-Peacock et al., 2012) and electroencephalography (LaRocque et al., 2013) and is easily disrupted by non-invasive brain stimulation (Zokaei et al., 2014) compared with non-attended items. These results suggest that the attended memory items are in the active state, whereas other items are in the latent state (LaRocque et al., 2014; Olivers et al., 2011). However, it remains unknown how easily/quickly we can retrieve memories in each state and how much information can be retained in the active state.

In this study, we addressed the above issues with a novel behavioral paradigm in humans. In brief, we found that when the memory load (i.e., the number of memorized color/letter pairs) was two, the reaction times (RTs) of the two memory items were similarly short. However, when the memory load was more than two, the RTs of non-recent or unattended memory items were sharply prolonged relative to the most recent or attended memory items, having behavioral implications for the dissociation between the active and latent states in working memory.

## MATERIALS AND METHODS

### Participant details

Fifteen participants (mean age ± standard deviation of 21.1 ± 2.0 years; 7 females, one left-handed) participated in the experiment under Load 2, 3, and 4 conditions and the Load 5 attending condition (see the Experimental design section for details of these conditions). Another 15 participants (mean age ± standard deviation of 23.8 ± 2.7 years; 7 females, one left-handed) participated under the Load 5 condition. The numbers of participants were set by referring to similar studies on the difference between attended and non-attended memory items in the working memory of humans (LaRocque et al., 2013; Lewis-Peacock et al., 2012; Zokaei et al., 2014). All participants had normal or corrected-to-normal visual acuity and reported no color vision deficiency. Participants gave informed consent before the experimental session and received monetary compensation. All experimental protocols and procedures were approved by the SFC Research Ethics Committee, Keio University (Approval Number 293).

### Experimental design

Throughout all experiments, the stimuli and program were generated using MATLAB (MathWorks, Natick, MA, USA) and the Psychophysics Toolbox (Psychtoolbox-3) (Brainard, 1997). A monitor was positioned approximately 57 cm from the participant. In each trial, a colored circle (having a visual angle of 1.5° × 1.5°) was presented at the center of the screen for 1500 ms, followed by a corresponding lowercase letter of the alphabet. The letter remained on the screen until a response was given by the participant. The eight candidate colors presented were [230,0,18], [243,152,0], [207,219,0], [34,172,56], [0,160,233], [29,32,136], [146,7,131], and [228,0,127] in the [R, G, B] color space. The corresponding letter candidates were s, d, f, j, k, and l. Within the same set, the color/letter pairs were consistent, and participants were asked to memorize the pairs. Once the memory was formed, the participants were instructed to respond quickly by giving the corresponding letter when the color was presented, before the presentation of the corresponding letter. They were asked to give the letter as quickly as possible with their ring, middle, and index fingers of both hands, using a keyboard. Feedback on correct or incorrect answers was displayed immediately after the participant’s response for 500 ms. In the case of a correct response, the correct letter was displayed in blue; in the case of an incorrect response, the correct letter was displayed in red. The inter-trial interval was set at 1000 ms, and during the interval, a fixation cross was displayed at the center of the screen. Under the Load N condition, the number of color/letter pairs was N. In each set of trials, each pair was presented 20 times, yielding 40 trials for the Load 2 condition, 60 trials for the Load 3 condition, 80 trials for the Load 4 condition, and 100 trials for the Load 5 condition. The number of sets was adjusted so that the number of trials for each load condition was at least 300 (eight sets for Load 2, five sets for Load 3, four sets for Load 4, and three sets for Load 5 conditions). Under the Load 5 attending condition, participants were required to internally attend the color/letter pair of the very first trial in each set to investigate the possibility of whether humans can intentionally change the state of a memory item. Under this condition, the inter-trial interval was set at 2000 ms to provide time to attend the memorized pair. Other settings were the same as those under the Load 5 condition.

### Data analyses

Under each condition, we separately calculated the RT distribution for successive and non-successive stimuli (Fig. 2a and Fig. 4b) and attended and non-attended stimuli (Fig. 4a). Successive trials are the trials whose stimulus is of the same color as the stimulus in the most recent trial whereas non-successive trials are the other trials. Attended trials are the trials whose stimulus is attended due to experimental instruction whereas non-attended trials are the other trials. Participants were required to answer within 1500 ms of the start of each trial; data with RTs over 1500 ms were removed. The distribution of RTs was calculated through kernel density estimation with a Gaussian kernel (bandwidth of 30 ms). Additionally, we analyzed the RT distribution depending on recency, which was classified according to how many trials ago the participant encountered the same stimulus (Fig. 3). For example, a recency of n = 2 means that the participant encountered the same stimulus two trials ago. RTs were analyzed from n = 1 to n = 5.

## Statistical analysis

In comparing data between two conditions, we conducted paired t-tests because normality for almost all the data for this comparison was verified in a Shapiro–Wilk test. For the data whose normality was not verified, we conducted both a paired t-test and Wilcoxon signed-rank test. In comparing more than three conditions, we conducted a one-way analysis of variance when the normality of the data was verified and a Kruskal–Wallis test when the normality was not verified.

## RESULTS

### Overflow of memories into a latent state

In the experiment, participants remembered color/letter pairs and answered the corresponding letter quickly when a colored circle appeared on the screen (Fig. 1a). Figure 1b shows the RT as a function of the number of encounters, revealing that the participants could give the corresponding letter in 1500 ms (before the appearance of the answer) after the second encounter and confirming that they successfully remembered the color/letter pairs in the first encounter. Here, we divided the stimuli into successive and non-successive stimuli (see, Fig. 1a). As a result, we found that as the memory load increased, the RTs for non-successive stimuli gradually separated from those for successive stimuli (Fig. 2a). A comparison of the average RTs across participants (Fig. 2b) showed that RTs for successive stimuli were significantly shorter than those for non-successive stimuli under the Load 3 condition (successive, 493.8 [95% confidence interval (CI), 464.5–523.2] ms; non-successive, 553.0 [95% CI, 499.4–606.7] ms; t15 = −3.52, p < 0.0034), Load 4 condition (successive, 520.4 [95% CI, 493.9–547.0] ms; non-successive, 654.9 [95% CI, 601.6–708.1] ms; t15 = −7.88, p < 0.001), and Load 5 condition (successive, 543.9 [95% CI, 520.2–567.6] ms; non-successive, 763.6 [95% CI, 711.3-816.0] ms; t15 = −11.40, p < 0.001) conditions but not under the Load 2 condition (successive, 420.6 [95% CI, 392.7–448.5] ms; non-successive, 416.2 [95%CI, 389.4–443.0] ms; t15 = −0.62, p = 0.5421). Under the Load 3 condition, where the normality of RTs for non-successive stimuli was not verified with the Shapiro–Wilk test, we also applied the Wilcoxon sign-rank test, confirming the significant differences in RTs between successive and non-successive stimuli (p < 0.001).

**Fig. 1.**
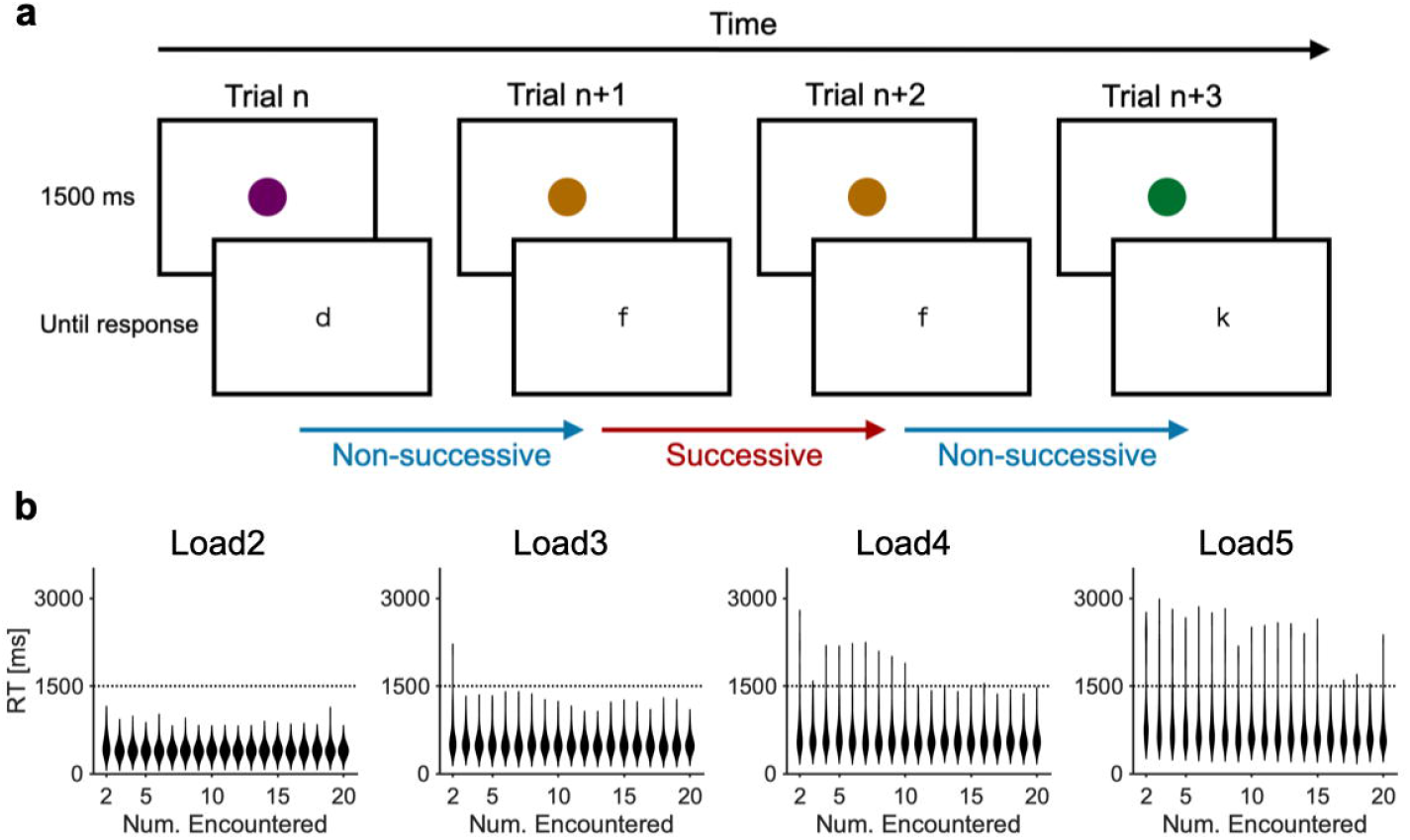
Task procedure. **a**, Experimental protocol. According to memorized color/letter pairs, participants quickly pressed the corresponding key upon the appearance of a colored circle. The number of pairs to remember is referred to as the memory load. **b**, Distributions of RTs as a function of the number of encounters. This indicates that after the first encounter with the color/letter pair, participants could remember the corresponding memory under each load condition.

Additionally, the RT differences between successive and non-successive stimuli showed a significant increase as the memory load increased (Fig. 2c, one-way analysis of variance, F = 37.56, p < 0.001; Tukey–Kramer’s test, p < 0.05 for all pairs). These data clearly demonstrate the overflow of working memory items from an easily accessible state to a somewhat latent, non-prioritized state, with the capacity of the former state being two chunks of information.

### Only the most recent memory is in the active state

We next examined whether only the most recent memory is in a privileged active state and other memories are equally in a latent state, or whether accessibility decreases as a function of the recency of the memory. The RT distribution (Fig. 3a) shows that only the most recent memory had a uniquely fast RT under the Load 4 and 5 conditions. Statistically analyzing the RTs across participants, we confirmed the uniqueness of the most recent memory (Fig. 3b). Under the Load 2 and 3 conditions, the RTs across participants were not significantly different according to the recency (Kruskal–Wallis test, p = 0.43 under the Load 2 condition; p = 0.18 under the Load 3 condition). Under the Load 4 condition, the RTs across participants were significantly different according to the recency (Kruskal–Wallis test, p = 0.0039). Multiple comparisons under the Load 4 condition showed that RTs for recency of n = 1 were significantly shorter than those for n = 2, 3, 4 (p = 0.0076, 0.0079, and 0.0452, respectively) and marginally significantly shorter than those for n = 5 (p = 0.0507). Other pairs showed no significant differences. Under the Load 5 condition, the RTs across participants were significantly different according to the recency (Kruskal–Wallis test, p < 0.001). Multiple comparisons under the Load 5 condition showed that RTs for recency n = 1 were significantly shorter than those for other recency values (p < 0.001 for all pairs) and other pairs showed no significant difference (p > 0.05). These results indicate that accessibility to the most recent memory is distinctly high, whereas accessibility to the non-recent memories is uniform irrespective of the recency of the stimuli.

**Fig. 2.**
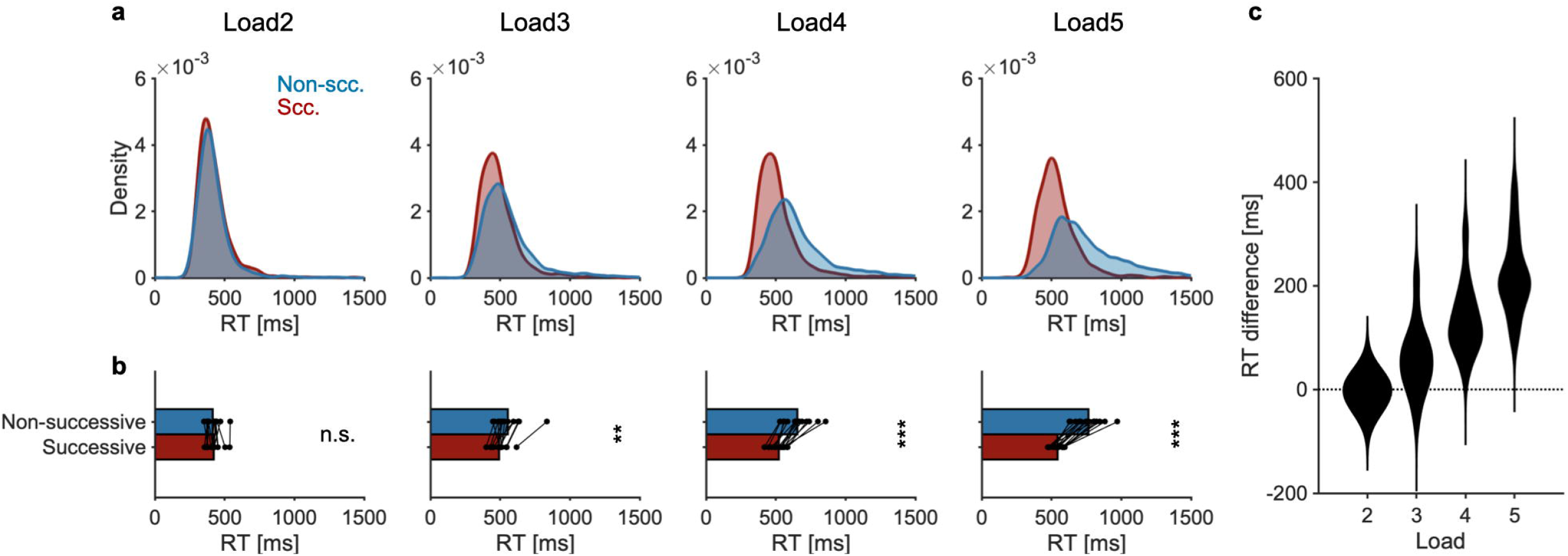
Overflow of memories from the easily accessible state. **a**, Grand-averaged RT distributions for successive (Scc.) and non-successive (Non-scc.) stimuli. **b**, RT comparison between successive and non-successive stimuli. Each dot denotes a participant. **c**, Distributions of the RT difference between successive and non-successive stimuli for each condition. As the memory load increased, RTs to of successive stimuli diverge from those of successive stimuli. n.s., not significant. **p<0.01, ***p<0.001.

**Fig. 3.**
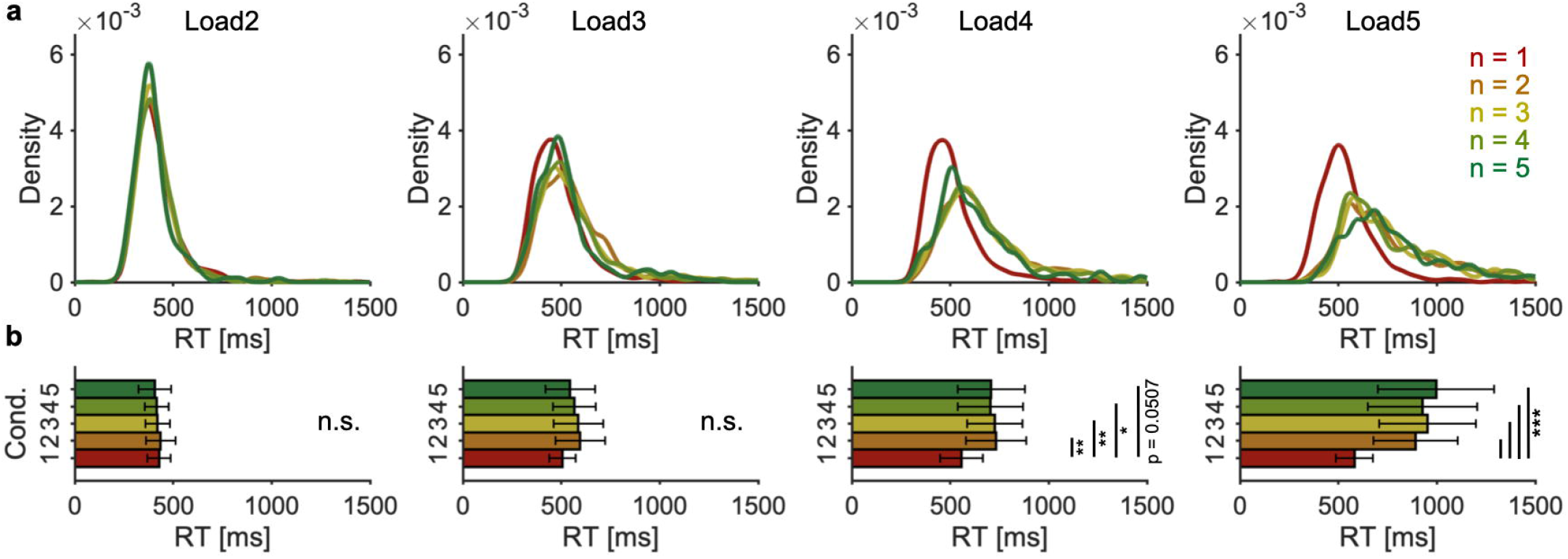
Relationship between RT distributions and the recency of memories. **a**, Grand-averaged RT distributions for each recency condition. Recency is classified according to how many trials ago the participant encountered the stimulus. **b**, Bar graphs showing average RTs across conditions. RTs of only the most recent memories are significantly shorter than others. n.s., not significant. *p < 0.05, **p < 0.01, ***p < 0.001.

**Fig. 4.**
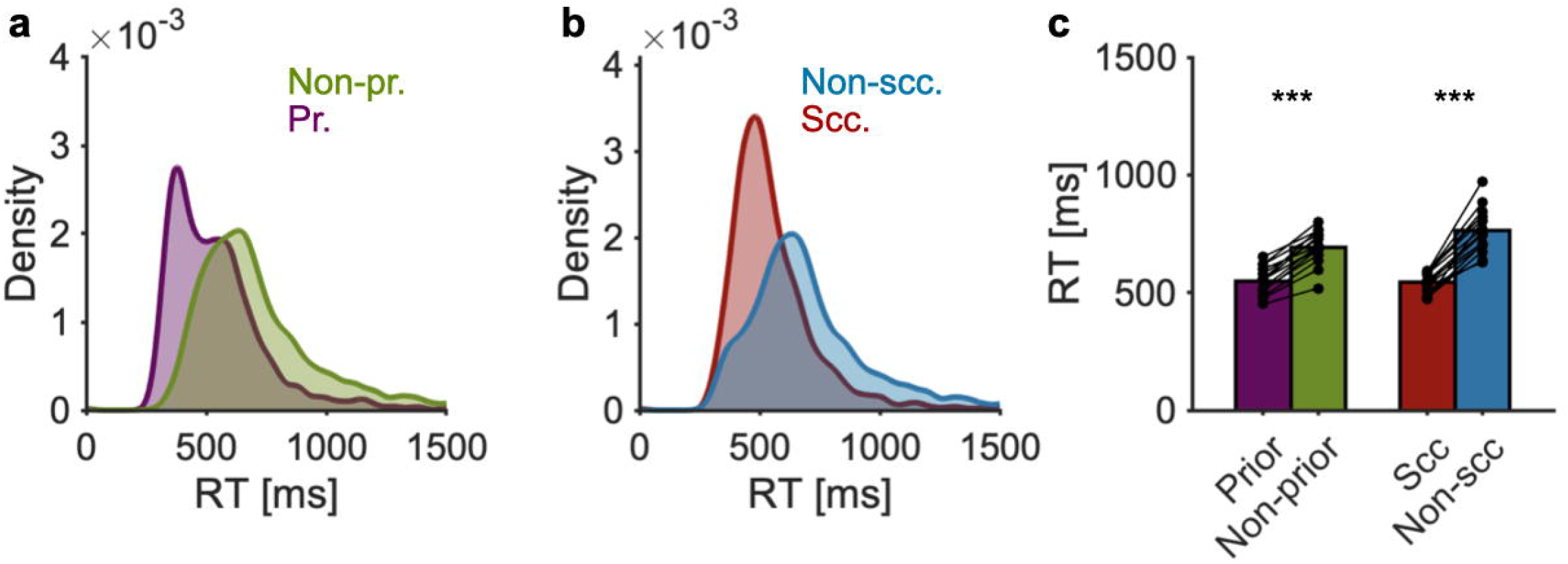
RT distributions for consciously prioritized memories. In an additional experiment, participants intentionally prioritized the color/letter pair that they encountered in the very first trial. **a**, Distributions of RTs for the prioritized (Pr.) and other (Non-pr.) stimuli (left) and those of successive (Scc.) and non-successive (Non-scc) stimuli (right). **b**, Bar graphs denoting individual RTs for each condition. Intentionally prioritized items are recalled more quickly than other items. ***p < 0.001.

### Internally attended memory enters the active state

By conducting an additional experiment, we examined whether internally attended memory enters the active state. It was found that the RTs of attended memory items were significantly shorter than those of non-attended items (attended, 550.0 [95% CI, 517.8–582.3] ms; non-attended, 693.6 [95% CI, 653.8–733.4] ms; t15 = −14.3731, p < 0.001). Additionally, the RTs of internally attended items were distributed similarly to those of the successive stimuli under the Load 5 condition of the previous experiment. The average RTs were not significantly different between the attended and successive stimuli (attended, 550.0 [95% CI, 517.8–582.3] ms; successive, 543.9 [95% CI, 520.2–567.6] ms; t15 = 0.3205, p = 0.7533). These results demonstrate that internally attended memories enter the active state and it is not critical when participants encounter them.

## DISCUSSION

The present study presented the behavioral characteristics of coexisting active and latent states in working memory. When the memory load was two, participants could recall both pairs similarly quickly irrespective of the recency of the memory. However, when the memory load was more than two, the RTs of the recall of the non-recent pairs were prolonged relative to those of the most recent pairs. The RTs of the recall of the non-recent memory became gradually prolonged as the memory load increased from two to five, as if the memory items overflowed from the easily accessible active state to the latent state. Interestingly, the recency effect of RTs was limited to the most recent memory item, whereas there was no significant difference in RTs among other memory items irrespective of how many trials ago the memory item appeared. In an additional experiment, similar to the result for the most recent memory items, we observed that the RTs of attended memory items were shorter than those of non-attended memory items.

Our results support the co-existence of active and latent states in working memory and provided the behavioral characteristics of each state. For a long time, working memory items have been believed to be maintained by the sustained firings of neurons, especially in the prefrontal cortex (Constantinidis et al., 2018; Funahashi et al., 1989; Fuster & Alexander, 1971). On the contrary, later studies showed that working memory items can be recalled after the extinction of the sustained activity of neurons (Rose et al., 2016; Spaak et al., 2017). These reports suggested that working memory items can also be maintained in the activity-silent, latent state without sustained firings of neurons (Stokes, 2015), possibly via short-term synaptic plasticity (Mongillo et al., 2008; Zucker & Regehr, 2002). Combining the results, it has been suggested that there are co-existing active and latent states in working memory (Kamiński & Rutishauser, 2020; LaRocque et al., 2014), and experimental support has been obtained in neuroimaging (LaRocque et al., 2013; Lewis-Peacock et al., 2012) and neurostimulation (Zokaei et al., 2014) paradigms. Here, we presented unquestionable RT differences between the putative active and latent states, adding strong evidence of the co-existence of the multiple states in working memory. Overall, our results indicate that dissociation in the neural states is robust enough to be observed through our behavior.

Importantly, our results demonstrate that the capacity of the active state in working memory is approximately two chunks of information. This finding is based on the observation that participants could recall both memories similarly quickly when the memory load was two. The “active memory capacity” of two is critically small compared with the traditionally considered working memory capacity of four (Cowan, 2001) or seven (Miller, 1956). Thus, not all items in the working memory capacity are uniform and there should be a dissociation of the capacity between the active state and the other state. Interestingly, the active memory capacity of two suggests that the active state of working memory is not equal to that of the conscious (Baars & Franklin, 2003; Trübutschek et al., 2019) or attended part of working memory (LaRocque et al., 2014; Olivers et al., 2011), whose capacity would be one at a time (Blake & Logothetis, 2002; Landau & Fries, 2012; Tong et al., 2006). Considering that the working memory capacity frequently declines in patients with developmental disorders (Alloway & Archibald, 2009) and the elderly (Brockmole & Logie, 2013) and is positively correlated with academic achievement (Jurden, 1995), we believe it is meaningful to examine how each capacity (active or other) is related to the above-mentioned cognitive differences. For instance, it would be possible that a certain disability diminishes only the capacity of the active state, whereas another type of disorder impairs the capacity of the latent state, but not the one of the active state.

The observation of RT prolongation as a function of the memory load has detailed implications for the latent state of working memory (Stokes, 2015; Wolff et al., 2017). The RTs of non-recent memory items were prolonged as the memory load increased relative to those of the most recent items. This prolongation of RTs indicates that the retrieval process of memory items in the latent state becomes increasingly difficult as the memory load increases. On the one hand, this prolongation would be explained by the “serial search” account of working memory retrieval (Sternberg, 1966, 2016), in which memory items are serially scanned one by one and matched items are then retrieved, analogous to a visual search (Kong & Fougnie, 2019; Wolfe, 2020; Wolfe & Horowitz, 2017). When we search for a visual target among distractors, the time required to find the target is prolonged in proportion to the number of visual stimuli, indicating the search is done by scanning the stimuli one by one (Moran et al., 2016; Treisman & Gelade, 1980). This is consistent with a recent study providing commonality between working memory and a visual search (Kong & Fougnie, 2019). On the other hand, our results are also consistent with the “decision-making” account of memory retrieval (Ratcliff, 1978). In this account, because memory items whose internal evidence (e.g., corresponding neural activities) reaches a threshold are retrieved, the RT is prolonged when the memory load is high (Pearson et al., 2014). Further work is needed to determine the exact process of working memory retrieval.

Regarding the recency effect, the results suggest that the most recent memory is in the active state and the other memories are in the latent state uniformly. This privilege of the most recent memory is consistent with previous reports on perceptual bias toward the most recent stimulus (Akrami et al., 2018; Barbosa et al., 2020; Fischer & Whitney, 2014; Murai & Whitney, 2021; Raviv et al., 2012). That is to say, new sensory input is perceived or memorized in a biased location toward the most recent trial. Additionally, the most recent memory has been found to be fragile through external brain stimulation (Zokaei et al., 2014). Combining these results with our results suggests that the most recent memory is special in working memory and easily accessible in recalling compared with non-recent memories.

In conclusion, our results demonstrated that the most recent memories and the internally attended memories are in the easily accessible active state and recalled more quickly than other memories. Importantly, we found that an active memory capacity of two differs from that of other cognitive capacities, such as the traditionally considered working memory capacity of four to seven and consciousness and attention capacity of one. In future work, elucidating the corresponding neural mechanisms will be one of the most urgent issues. Additionally, it will be crucial to classify the traditionally considered working memory into the active memory capacity and the latent memory capacity. We should then re-examine relationship of these capacities with our daily activity and clinical issues.

## Acknowledgment

This work was supported by a designated donation from Living Platform, Ltd, Japan to J.U., Taikichiro Mori Memorial Research Grants to T.I., Sasakawa Scientific Research Grant (The Japan Science Society, JSS) to T.I., and JST SPRING, Grant Number JPMJSP2123 to T.I. We thank Ms. Tomomi Hamaoka, Ms. Kana Iijima, and Ms. hieko Matsuda for their secretarial assistance and all other members of our laboratory for their insightful comments on the work. We thank Edanz (https://jp.edanz.com/ac) for editing a draft of this manuscript.

